# The protein-tyrosine phosphatase Shp2 is essential for lymphatic endothelial cell differentiation in zebrafish

**DOI:** 10.1101/2025.09.09.675100

**Authors:** Daniëlle T.J. Woutersen, Andreas van Impel, Stefan Schulte-Merker, Jeroen den Hertog

## Abstract

Mutations in genes associated with the VEGFC/VEGFR3 signaling pathway are strongly linked to a range of developmental and functional defects in the lymphatic system. *PTPN11* encodes SHP2, a protein-tyrosine phosphatase that plays a critical role in receptor protein-tyrosine kinase (RTK) signaling. In this study, we investigated lymphangiogenesis in mutant zebrafish embryos lacking functional Shp2 and demonstrate that Shp2 is essential for normal lymphatic development in the trunk and head regions of zebrafish embryos. Embryos lacking functional Shp2 failed to develop lymphatic vessels and lymphatic endothelial cells (LECs), whereas venous intersegmental vessel (vISV) formation remained unaffected. While endothelial cells sprouted normally from the posterior cardinal vein (PCV), they did not migrate to the horizontal myoseptum to form parachordal lymphangioblast (PL) cells. Rescue experiments using Shp2 mutants demonstrated that catalytic activity and signaling properties of Shp2 are required for restoration of normal lymphangiogenesis. These findings highlight a pivotal role for Shp2 signaling in the migration and differentiation of lymphatic endothelial cells without affecting venous angiogenesis.

## Introduction

The lymphatic vasculature is comprised of a specialized network of capillaries and vessels dedicated to maintain fluid homeostasis, to facilitate immune surveillance and to reabsorb dietary fat. A functional lymphatic vasculature is a prerequisite to prevent pathophysiological conditions including lymphedema and inflammation. Over the years, substantial discoveries regarding the development of the lymphatic vasculature have been made using the zebrafish (*Danio rerio*). The quick ex-utero development, translucency of the embryos and the availability of fluorescent transgenic labels provide a unique opportunity to visualize lymphatic vessel formation over time (Castranova et al., 2021; Das et al., 2022; Hogan, Bos, et al., 2009; Hogan, Herpers, et al., 2009; Hogan & Schulte-Merker, 2017; Klems et al., 2020; Wang et al., 2020).

Lymphangiogenesis, the process of lymphatic vessel formation by sprouting, is initiated in the zebrafish trunk after completion of primary angiogenesis. Venous endothelial cells from the posterior cardinal vein (PCV) start to divide and form sprouts from the PCV towards neighboring intersegmental vessels. Next, endothelial sprouts protrude dorsally at around 32 hours post fertilization (hpf) (Bussmann et al., 2010). Once these secondary sprouts encounter an arterial intersegmental vessel (aISV), roughly half of these sprouts establish a stable connection and adopt the venous identity (Geudens et al., 2019). Subsequently, these venous intersegmental vessels (vISVs) become perfused. The secondary sprouts that do not stably connect with an intersegmental vessel (ISV) migrate further into the direction of the horizontal myoseptum (HM). Once these cells reach the HM they are referred to as parachordal lymphangioblast (PL) cells (Hogan, Bos et al., 2009). At 72 hpf the PL cells start dividing and migrate in dorsal and ventral direction along the aISVs to constitute the intersegmental lymphatic vessels (ISLVs), the thoracic duct (TD) and the dorsal longitudinal lymphatic vessel (DLLV)(Cha et al., 2012; Bussmann et al., 2010; Geudens et al., 2019). The TD and DLLV are interconnected via the ISLVs aligning with the aISVs, thus forming the lymphatic vasculature of the embryonic trunk, which is completed at 5 dpf (Bussmann et al., 2010).

Whereas trunk lymphangiogenesis has been described extensively, development of lymphatic structures in the head has been highlighted only recently (Eng et al., 2019; Okuda et al., 2012). The trunk lymphatics are exclusively derived from venous origin, yet lineage-tracing experiments indicate that the lymphatic structures in the head comprise both venous and non-venous derived progenitor populations (Eng et al., 2019). The common cardinal vein (CCV) gives rise to the facial lymphatic sprout (FLS), which is complemented with cells from the primary head sinus (PHS). Eventually, a third population of ventral aorta lymphangioblasts (VA-L) is added sequentially to the tip of the FLS (Eng et al., 2019).

The vascular endothelial growth factor-C (VEGFC)/ VEGF Receptor-3 (VEGFR3) signaling pathway is essential for lymphangiogenesis in the trunk (Hogan, Bos, et al., 2009; Hogan, Herpers, et al., 2009; Le Guen et al., 2014; Shin et al., 2016; Villefranc et al., 2013). Various mutant zebrafish models have been generated illustrating the importance of this signaling pathway in lymphangiogenesis. Interestingly, genetic alterations in genes associated with this pathway are strongly linked to congenital lymphatic disease in human patients. Particularly, next to VEGFC and VEGFR3, extracellular factors involved in activation of the secreted VEGFC protein, such as ADAMTS3 and CCBE-1 (Hogan, Bos, et al., 2009; Wang et al., 2020), as well as its receptor, VEGFR3, are implicated in the process (Hogan, Bos, et al., 2009). Yet, the intracellular signaling mediators that act downstream of the VEGFR3 receptor protein-tyrosine kinase in lymphatic endothelial cells (LECs) are less well characterized.

We are interested in the function of the ubiquitously expressed cytoplasmic Src homology region 2 (SH2)-containing protein tyrosine phosphatase 2 (SHP2). SHP2 is broadly expressed and has a central role in receptor protein-tyrosine kinase (RTK) signaling (Feng et al., 1993; Freeman et al., 1992; Vogel et al., 1993). SHP2 is indispensable during embryonic development as shown in both mice and zebrafish (Bonetti et al., 2014; Yang et al., 2006). As a result of teleost-specific whole genome duplication, zebrafish contain two paralogs of the protein-tyrosine phosphatase non-receptor type 11 (*ptpn11)* gene encoding the functionally redundant Shp2a and Shp2b proteins. Zebrafish double mutants lacking functional Shp2 altogether display a pleiotropic phenotype, including mild to severe edemas (Bonetti et al., 2014). Hence, we were interested to investigate lymphangiogenesis in zebrafish embryos lacking functional Shp2.

Here we report that Shp2 is essential for normal lymphangiogenesis in zebrafish. Embryos lacking functional Shp2 do not form lymphatic structures. Whereas endothelial cells from the PCV sprout normally, they do not migrate to the horizontal myoseptum to form PL cells. Instead, these sprouts retract and sometimes disintegrate during the process. Interestingly, vISVs form normally. We conclude that Shp2-mediated signaling is essential for lymphangiogenesis.

## Materials and Methods

### Zebrafish husbandry

All procedures involving experimental animals were approved by the animal experiments committee of the Royal Netherlands Academy of Arts and Sciences (KNAW), Dierexperimenten commissie protocol HI18-0702 and HIdHe16596.22.01, and conducted according to local guidelines in compliance with national and European law. Zebrafish were maintained under standard laboratory conditions (Aleström et al., 2020) and staged as previously described (Kimmel et al., 1995). The *ptpn11a^hu3459^* and *ptpn11b^hu5920^* knockout lines were previously generated (Bonetti et al., 2014). Published transgenic lines used in this study were *Tg(fli1a:eGFP)^y1^* (Lawson & Weinstein, 2002), *Tg(kdrl:*mCherry-CAAX)*^y171^* (Fujita et al., 2011), *Tg(flt1:tdTomato)^hu5333^* (Bussmann et al., 2010) and *Tg(flt4:*mCitrine*)^hu7135^* (van Impel et al., 2014). Fertilized eggs were incubated at 28.5°C in E3 medium (5 mM NaCl, 0.17 mM KCL, 0.33 mM CaCl_2_, 0,33 mM MgSO_4_) and occasionally pigmentation was prevented by adding phenylthiourea (Sigma-Aldrich, P7629) at a concentration of 0.003% (v/v) to the E3 medium at 24 hpf.

### Genotyping

All zebrafish in this study were genotyped to establish the *ptpn11a* and *ptpn11b* genotype. Genomic DNA was extracted by lysis of whole embryos or fin clip tissue as described (Hale & den Hertog, 2018). Genotyping was performed by Sanger sequencing (Macrogen Europe B.V., Amsterdam, The Netherlands) of gene-specific PCR products amplified with primers for *ptpn11a* (exon 3) 5’-GCGCTGTCACACACATTAAGA and 5’-TCCCAAATTGTCATGTAAGG and *ptpn11b* (exon 3) 5’-GTCTGTCATCCCTCATTTCC and 5’-GCAGGATTTATTCTGTCCAC.

### Constructs, mRNA synthesis and microinjections

All mRNA used in this study was synthesized from pCS2+ plasmids containing an N-terminal eGFP followed by a peptide-2A cleavage sequence and Shp2a or Shp2b. Point mutations were introduced by Q5 site-directed mutagenesis (New England Biolabs, E0554S). 5’ capped sense mRNA was synthesized from Not1 linearized constructs using the mMessage mMachine SP6 kit (Ambion, AM1340). Embryos were injected at the one-cell stage with 1 nL of the mRNA solution as previously described (Hale & den Hertog, 2016).

### Brightfield microscopy, confocal microscopy and time-lapse imaging

Brightfield microscopy was performed with a Leica M165 FC stereomicroscope (Leica Microsystems Wetzlar, Germany) connected to a Leica DMC5400 camera (Leica Microsystems Wetzlar, Germany). Confocal microscopy and time-lapse imaging were performed with a Leica SP8 (Leica Microsystems Wetzlar, Germany). Fixed embryos were mounted in 2% methylcellulose on a microscope slide with a concave depression cavity. Live embryos were anesthetized with 0.1% tricaine methane sulfonate, mounted on glass cover slips with 0.6% UltraPure agarose and covered with E3 medium. Confocal microscopy and time-lapse imaging were captured using a 10x or 20x objective and z-stack step size ranging between 1.5 and 2.0 µm. All images were examined using Imaris and/or ImageJ software (US National Institutes of Health, Bethesda, MD).

### Whole mount *in situ* hybridization

PTU treated embryos were euthanized in 50% tricaine methane sulfonate and fixed in 4% paraformaldehyde at 4°C. Whole mount *in situ* hybridization was performed as previously described (Thisse & Thisse, 2008). Digoxigenin-labelled probe synthesis was performed using plasmids.

### Quantitative PCR

For quantitative PCR, 5 dpf embryos were genotyped by dissection of part of the tail, which was used for genotyping. Five wild-type and five *ptpn11a^−/−^ptpn11b^−/−^*embryos were selected and pooled per genotype, RNA was extracted and cDNA was made. Quantitative real-time PCR was performed using FastStart Universal SYBR Green Master Mix on a Bio-Rad CFX384 Touch Real-Time PCR Detection System. For each condition two biological replicates were used and each gene was measured in triplicate. The housekeeping gene beta-actin was used as internal reference. Analysis was performed using the Bio-Rad CFX Maestro software. To determine the fold difference in gene expression, the ΔΔCt method was used.

### Quantification and Statistical analysis

Statistics were performed in GraphPad Prism 8 (GraphPad Software, San Diego, USA version 8.2.1). For statistical analysis, embryos were blindly scored on the completeness of the TD, i.e. prior to genotyping. Absence of TD was scored as 0, partial presence of TD as 1 and complete presence of TD was included as 2. Statistical tests included Standard t-test and Mann-Whitney U test as indicated. All microinjection experiments were performed at least in triplicate, unless stated otherwise.

## Results

### Impaired development of the lymphatic vasculature in zebrafish embryos that lack functional Shp2

Previously, we used target-selected gene inactivation of both *ptpn11* paralogs to generate a stable knockout zebrafish line to study embryonic development in the absence of functional Shp2 (Bonetti et al., 2014). Loss of functional Shp2a in *ptpn11a^−/−^ptpn11b^+/+^* embryos induced a pleiotropic phenotype from 4 days post fertilization (dpf) onwards (**Figure 1a-d**) and was previously shown to be embryonic lethal (Bonetti et al., 2014). *Ptpn11a^+/+^ptpn11b^−/−^*embryos did not display detectable morphological defects (**Figure 1e,f**) and adult fish lacking functional Shp2b are viable and fertile (Bonetti et al., 2014). Double mutants (*ptpn11a^−/−^ptpn11b^−/−^*) developed severe morphological defects with eye, pericardial and yolk sac edemas at 5 dpf (**Figure 1g,h**). The defects in double mutants were more severe than the defects observed in *ptpn11a^−/−^ptpn11b^+/+^* embryos (*cf.* **Figure 1c,d and 1g,h**) and double mutants were embryonic lethal. Taken together, loss of Shp2 function induced severe edemas in zebrafish embryos.

**Figure 1.**
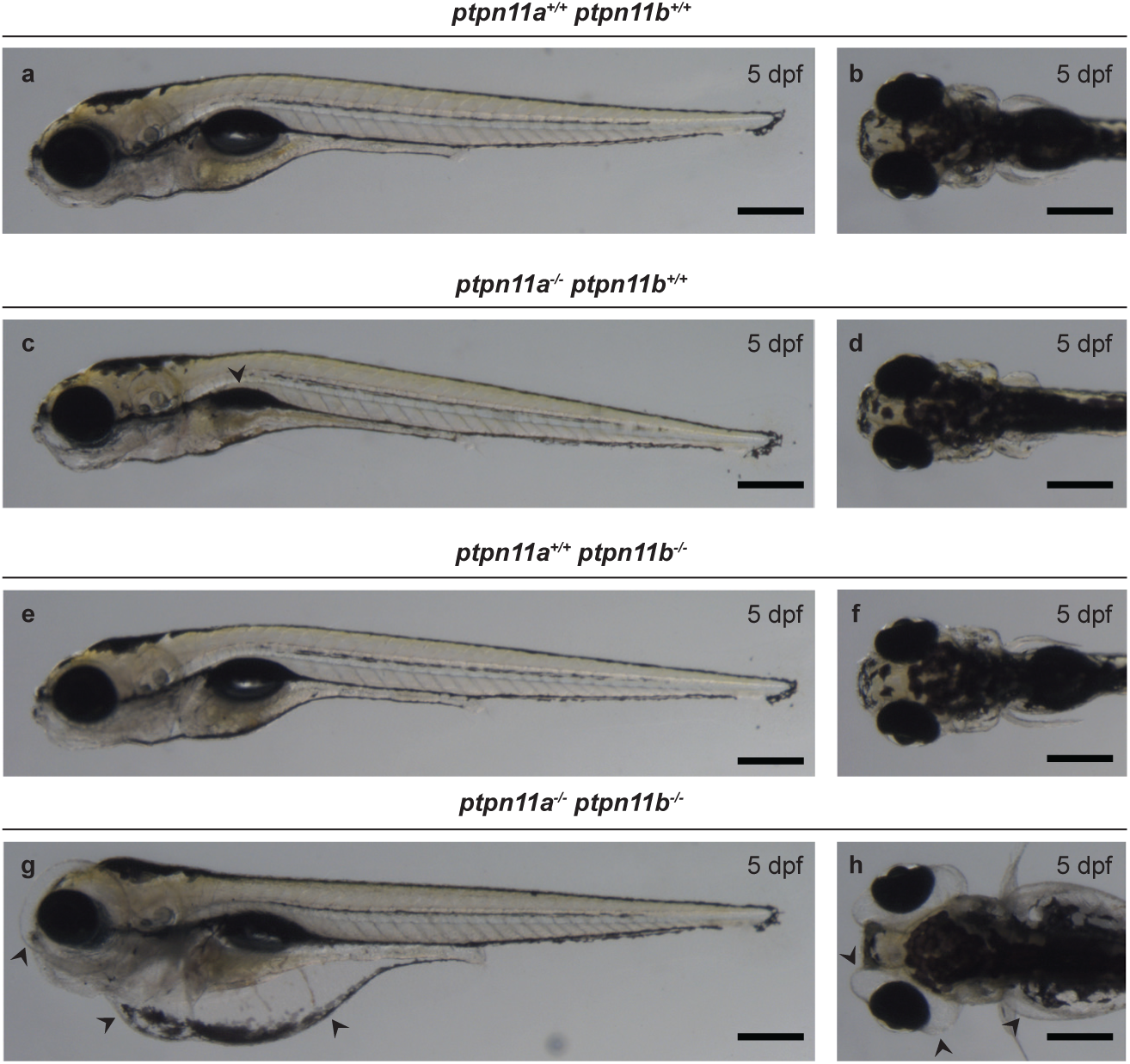
Lymphedema in double homozygous Shp2 knockout embryos. (**a-h**) *Ptpn11a^+/−^ ptpn11b^+/−^* fish were incrossed and representative brightfield images of 5 dpf zebrafish embryos of selected genotypes as indicated in lateral and dorsal orientation are shown. Ablation of the Shp2a protein yields a slim phenotype with an underdeveloped swimbladder (arrowhead) (**c-d**), whereas knockout of Shp2b does not yield a morphological phenotype (**e-f**). Double homozygous mutants display severe lymphedema including eye, cardiac and abdominal edema (**g-h**). Arrowheads indicate the edematous areas and underdeveloped swimbladder. Scalebar indicates 300 µm.

Severe edemas might be caused by compromised lymphatic function. To investigate whether there are defects in the lymphatic vasculature in the absence of Shp2, we imaged the lymphatic vasculature in double transgenic *flt4:mCitrine;flt1:tdTomato* embryos at 4 dpf. The *flt4:mCitrine* transgene highlights venous and lymphatic endothelial cells while *flt1:tdTomato* marks arterial endothelial cells (**Figure 2a-d**). Wild-type siblings and single mutants showed normal lymphatic vasculature whereas the *ptpn11a^−/−^ptpn11b^−/−^*mutant embryos lacked all lymphatic structures in the trunk including the ISLVs, DLLVs and the TD (**Figure 2d**).

**Figure 2.**
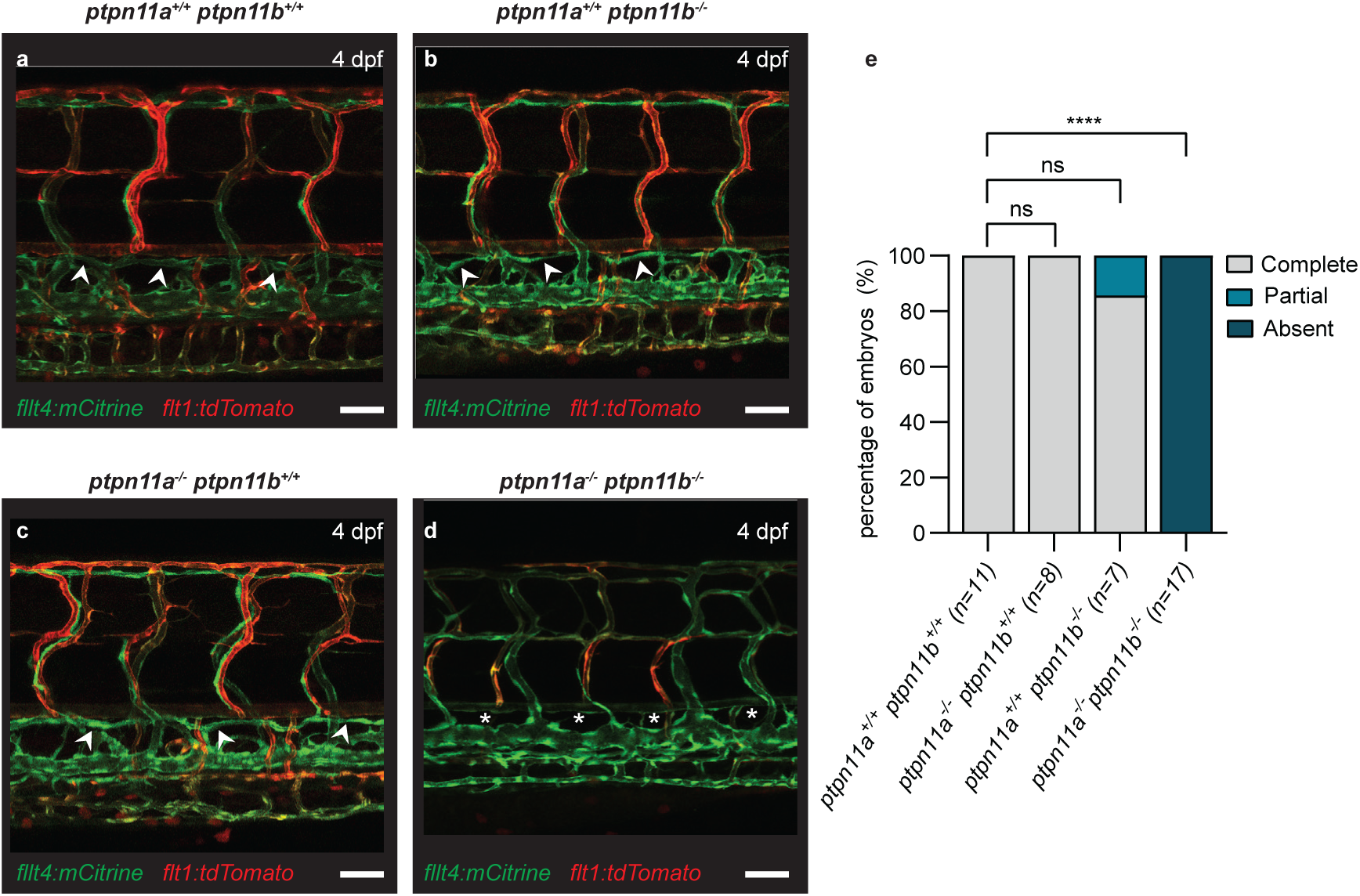
Shp2 is indispensible for ISLV, DLLV and TD formation in the zebrafish trunk. (**a-d**) *Ptpn11a^+/−^ptpn11b^+/−^* fish in *Tg(flt4:mCitrine;flt1:tdTomato)* background were incrossed and representative confocal projections of the trunk vasculature positive for flt4:mCitrine and flt1:tdTomato of 4 dpf wild-type, single and double homozygous mutants for Shp2 are shown. Scale bar indicates 50 µm. **(e)** Quantification of the percentage of embryos with a complete, partial or absent thoracic duct at 4 dpf. Statistical analysis was done using the Mann-Whitney U test. *p <0.05, **p <0.01, ***p <0.001, ****p <0.0001 and ns = not significant.

Some established zebrafish mutants, including *svep1^−/−^* and the *tabula rasa* mutant, which lacks functional Growth factor receptor bound protein 2b (Grb2b), have been reported to display variable penetrance with regard to the severity of the lymphatic phenotype (Hußmann et al., 2023; Karpanen et al., 2017; Mauri et al., 2021). Hence, we quantified the presence of the TD in the trunk at 5 dpf (**Figure 2e**). Embryos were scored in three categories: complete TD, partial TD or absent TD. In all homozygous double mutant embryos, the TD was completely absent at 5 dpf, which was significantly distinct from wild-type siblings (p<0.001). In fact, all lymphatic vessels were absent in the trunk. Whereas occasionally the TD was missing in a single segment of the single mutants, no significant lack of lymphatic structures was observed after loss of functional Shp2a or Shp2b alone, compared to wild-type siblings. These results indicate that Shp2a and Shp2b fulfill redundant roles during lymphatic development since lymphatic defects in the embryonic trunk are only evident after complete loss of functional Shp2. Therefore, we focused our analyses on the comparison between wild-type (*ptpn11a^+/+^ptpn11b^+/+^*) and double mutant (*ptpn11a^−/−^ptpn11b^−/−^*) embryos.

In addition to the trunk lymphatic vasculature, we examined lymphatic structures in the head (**Figure 3**). To this end, we assessed the appearance of the lymphatic vasculature in the head of transgenic *flt4:mCitrine ptpn11a^−/−^ptpn11b^−/−^* zebrafish embryos at 3 and 5 dpf and observed that whereas the lymphatic structures begin to form at 3 dpf, most of the facial lymphatic vasculature was missing compared to wild-type embryos at 5 dpf (**Figure 3**). Notably, the otolithic lymphatic vessel (OLV) and the medial facial lymphatic (MFL) vessel were missing in double mutant embryos at 5 dpf (**Figure 3b,e**). The lateral facial lymphatic (LFL) vessel appeared to be severely impaired in its development. Importantly, the facial collecting lymphatic vessel (FCLV), which forms the connection between facial lymphatics and the venous system (Shin et al., 2016) and whose development is governed by Svep1/Tie1 signaling independently of the Vegfc/Flt4 axis (Hußmann et al., 2023), still formed in *shp2* double mutants at 5 dpf (**Figure 3d-f**). Furthermore, another lymphatic cell population in the head region whose development depends on the Vegfc/Flt4 signaling axis, is a group of cells termed mural lymphatic endothelial cells, fluorescent granular epithelial cells or brain lymphatic endothelial cells (BLECs) (Bower et al., 2017; Galanternik et al., 2017; van Lessen et al., 2017). These cells populate the meningeal layer of the brain where they exert scavenging functions in wild-type embryos (Huisman et al., 2022). In *shp2* double mutants, however, this lymphatic cell population is absent at 5 dpf (*cf.* **Figure 3d,e**). Taken together, all lymphatic structures in the head, that depend on Vegfc/Flt4 signaling, were missing in *shp2* double mutant embryos at 5 dpf.

**Figure 3.**
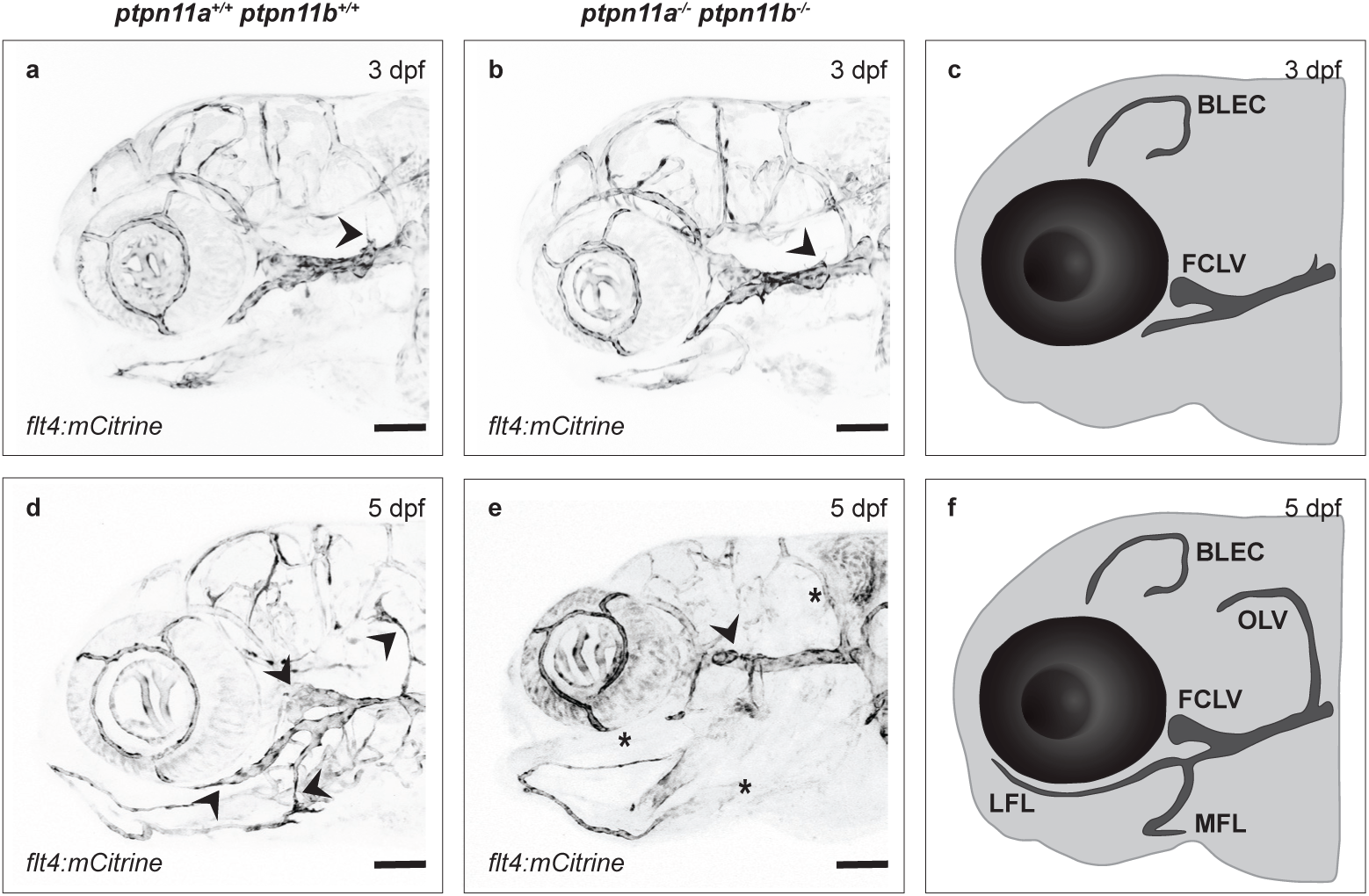
Lymphangioblast derived head lymphatic vessels are lacking in double homozygous mutants. Representative confocal projections are depicted of the lymphatics in the head staining positive in *Tg(flt4:mCitrine)* background at 3 and 5 dpf in (**a,d**) wild-type and (**b,e**) double homozygous mutants. Arrowheads indicate the presence and asterisk the absence of various facial lymphatic vessels. Scalebar indicates 50 µm. (**c,f**) Schematic representation highlighting the lymphatic vasculature at 3 and 5 dpf in wild-type embryos. BLEC, brain lymphatic endothelial cell; FCLV, facial collecting lymphatic vessel; OLV, otolithic lymphatic vessel; MFL, medial facial lymphatic; LFL, lateral facial lymphatic.

To verify that the lymphatic knockout phenotype observed in *ptpn11a^−/−^ptpn11b^−/−^*zebrafish embryos was caused by lack of functional Shp2, we performed rescue experiments. Synthetic mRNA encoding zebrafish Shp2a or Shp2b was micro-injected into offspring of an incross of *fli1a:eGFP* positive *ptpn11a^+/−^ptpn11b^−/−^* adults at the one-cell stage and the appearance of the TD was scored. In both *shp2a-* and *shp2b-*injected *ptpn11a^−/−^ptpn11b^−/−^*embryos the percentage of embryos with a partial or complete TD was significantly increased, compared to uninjected double mutants (**Figure 4**). Also, the edemas observed in the eye, cardiac and abdominal regions were rescued by the injections (**Supplementary Figure 1**). Microinjection of mRNA encoding Shp2a or Shp2b did not significantly affect TD development in wild-type embryos, suggesting that overexpression of Shp2 has no negative effects on lymphatic development. Taken together, the combined loss of Shp2a and Shp2b impaired development of the lymphatic vasculature in both head and trunk of the zebrafish embryo.

**Figure 4.**
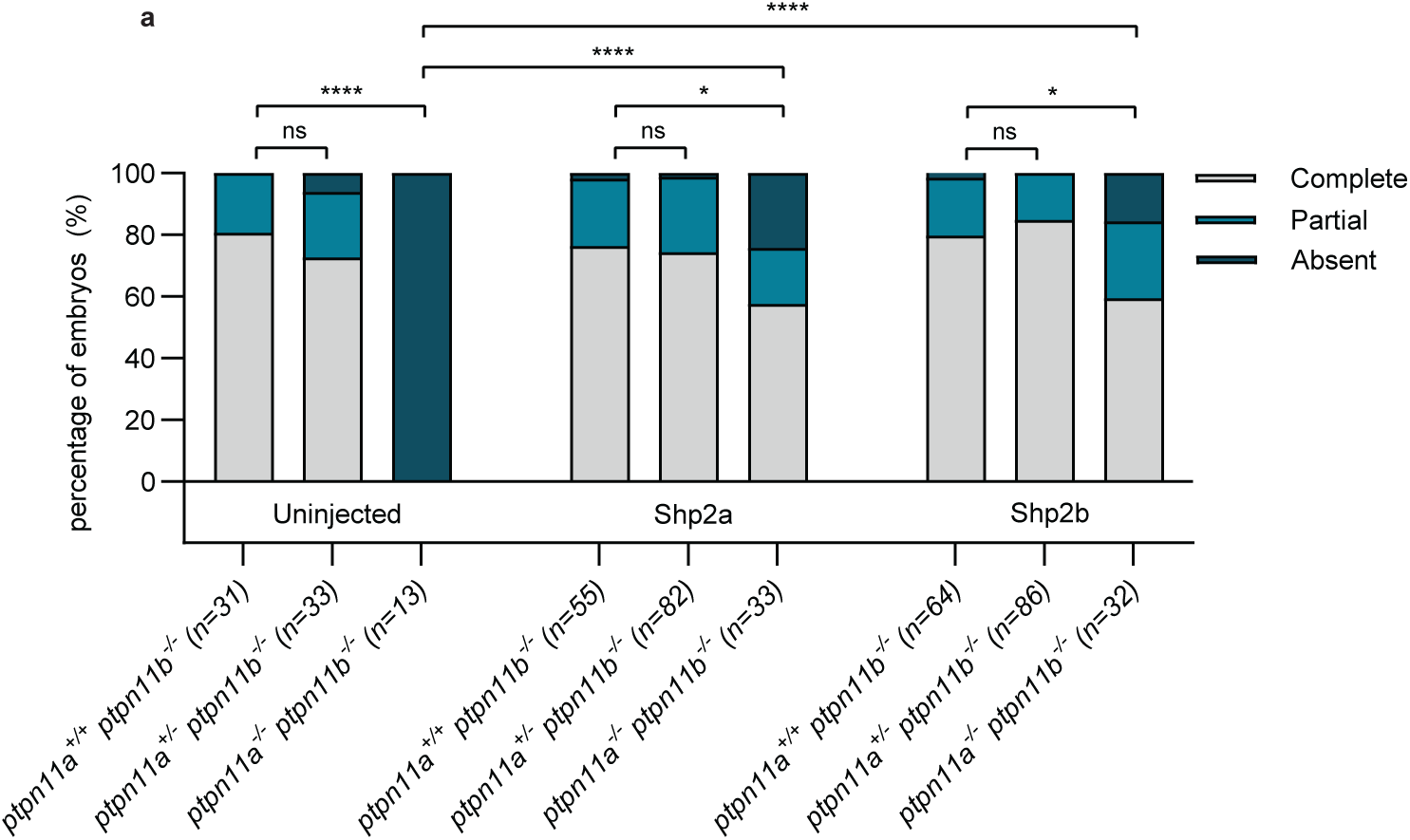
Shp2a and Shp2b expression rescues lymphangiogenesis defects in Shp2 knockout zebrafish embryos. *Ptpn11a^+/−^ptpn11b^−/−^* fish in *Tg(fli1a:eGFP;kdrl:mCherry)* background were incrossed. Embryos were injected at the one-cell stage with synthetic mRNA encoding Shp2a or Shp2b. At 5 dpf, the presence or (partial) absence of the *fli1a*-positive TD was quantified in uninjected controls and embryos injected with Shp2a or Shp2b by analysis of 10 segments of each embryo. The results of three independent experiments are pooled with the total number of embryos (n) indicated. Statistical analysis was done using the Mann-Whitney test *p < 0.05; **p < 0.01, ***p < 0.001, ****p <0.0001 and ns = not significant.

### Knockout of Shp2 impairs parachordal lymphangioblast formation

The trunk lymphatic vessels in the zebrafish are derived from PL cells. At 54 hpf, these lymphatic precursor cells align at the HM and serve as a pool of lymphatic precursors in the embryonic trunk. As the PL cells migrate away from the HM, they move along the aISVs to give rise to various lymphatic vessels including the ISLVs, DLLVs and TD (Hogan, Bos, et al., 2009; Yaniv et al., 2006). To examine whether the absence of the TD in double homozygous mutants is correlated with a defect in PL cell formation, we quantified the number of PL cells over the length of 10 somites in *flt4:mCit* transgenic embryos at 54 hpf (**Figure 5**). All double homozygous mutants lacked all PL cells at 54 hpf (**Figure 5d**). In sibling embryos, there was no significant difference in the number of PL cells detected between wild-type and single mutants (**Figure 5e**). The lack of both TD and PL cells in double homozygous mutant embryos emphasized the importance of *ptpn11* during early lymphangiogenesis and raised the question whether PL cells were even generated at all.

**Figure 5.**
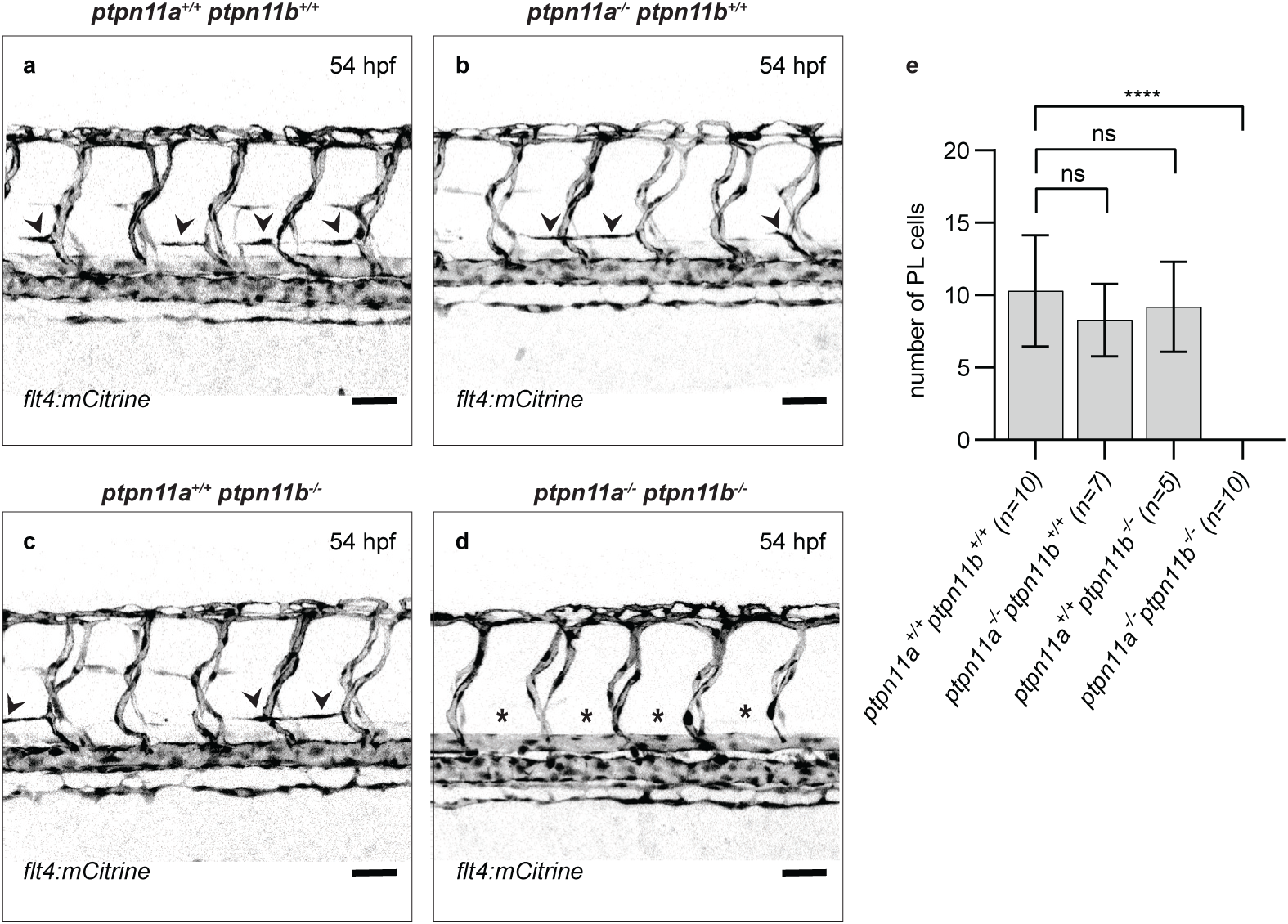
Parachordal lymphangioblasts are not formed in double homozygous mutants. (**a-d**) *Ptpn11a^+/−^ptpn11b^+/−^*fish in *Tg(flt4:mCitrine)* background were incrossed and representative confocal projections of the trunk vasculature of embryos at 54 hpf are shown of single and double homozygous mutants as indicated. Scalebar indicates 50 µm. (**e**) Quantification of the number of PL cells in the trunk at 54 hpf. Statistical analysis was done using the standard t-test *p < 0.05; **p < 0.01, ***p < 0.001, ****p <0.0001 and ns= not significant.

### Aberrant secondary sprouting in double homozygous mutants

We did not observe PL cells in double homozygous embryos lacking functional Shp2. This may be due to PL cells not forming at all, or to PL cells that formed, but disintegrated subsequently. To investigate this, we imaged the formation of PL cells by confocal time-lapse imaging of 8-10 adjacent intersegmental vessels in the *fli1a:eGFP* background between 30 and 70 hpf. PL cell formation starts with lympho-venous sprouting of endothelial cells from the PCV. In wild-type siblings, lympho-venous secondary sprouts emerged from the PCV and roughly half of these sprouts migrated towards the HM to become PL cells (**Figure 6a-f, supplemental movie 1**) as described before (Hogan, Bos, et al., 2009). In *ptpn11a^−/−^ptpn11b^−/−^* mutant embryos, sprouting initiated normally, but sprouts that did not connect to an aISV to generate an intersegmental vein, showed stalling behavior (**Figure 6i,j**), retracted back into the PCV or, in some cases, disintegrated (**Figure 6k-n, Supplementary Movie 2,3**). As a result, no sprouts migrated to the HM to form PL cells.

**Figure 6.**
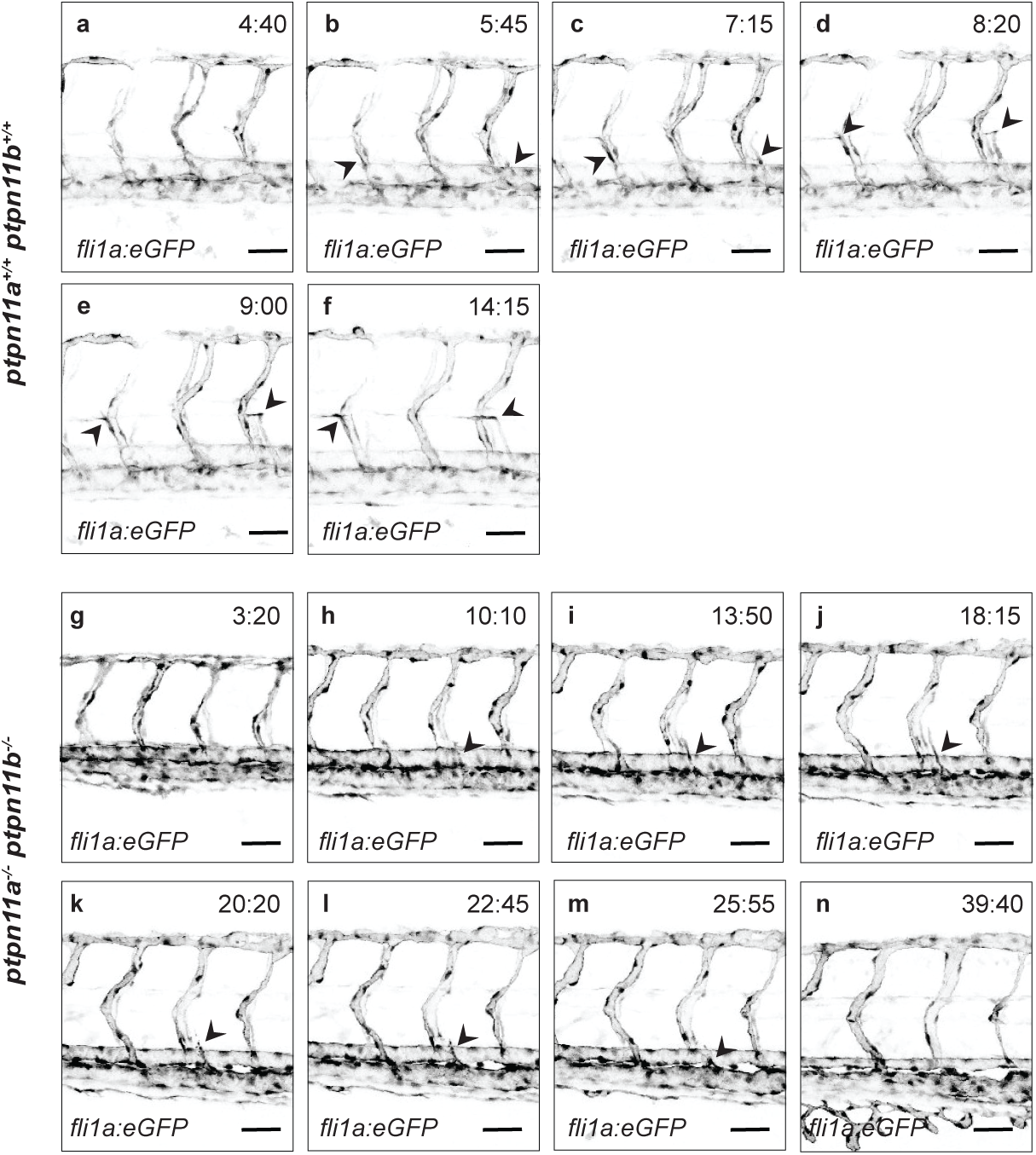
Aberrant secondary sprouting in double homozygous mutants. *Ptpn11a^+/−^ ptpn11b^−/−^*fish in *Tg(flt4:mCitrine)* background were incrossed and embryos were imaged by confocal microscopy. Cropped confocal projections are shown, extracted from time-lapse supplementary movies SM1 and SM2 of 8-10 adjacent ISVs during secondary sprouting events in (**a-f**) wild-type and (**g-n**) a representative double homozygous mutant embryo (t = 0:00 represents 32 hpf). Arrows indicate lymphatic sprouts emerging from the PCV. Scalebar indicates 50 µm.

The secondary sprouts that do not give rise to PL cells at the HM, connect stably to primary ISVs thereby remodeling them into vISVs (Bussmann et al., 2010; Geudens et al., 2019). Interestingly, formation of vISVs appeared to be unaffected in shp2 double mutant embryos (**Figure 6, Supplementary Movies 1-3**) as sprouts from the PCV connected with an aISV and formed a vISV, which became perfused after the connection was made. The formation of vISVs was indistinguishable between wild-type and double mutant embryos.

### Loss of Shp2 does not affect the arterial-venous ISV ratio

Normally, arteriovenous patterning has the tendency to alternate in the zebrafish trunk, yielding an aISV/vISV ratio of 1:1 (Bussmann et al., 2010). To examine if lack of functional Shp2 affects the arterio-venous patterning of the zebrafish trunk vasculature, we determined the number of aISVs and vISVs between segmental position 6 and 15 in *flt4:mCit;flt1:tdTomato* double transgenic embryos at 4 dpf (**Figure 7 a-d**). There was no significant difference in the aISV/vISV ratio between double homozygous mutant embryos and their siblings (**Figure 7e**). In addition, we observed normal perfusion and blood flow in the established vISV connections of the *ptpn11a^−/−^ptpn11b^−/−^* embryos and their siblings, suggesting that Shp2 is dispensable for the formation of vISVs. These results indicate that Shp2 is essential for early lymphangiogenesis in the trunk, but not for venous angiogenesis.

**Figure 7.**
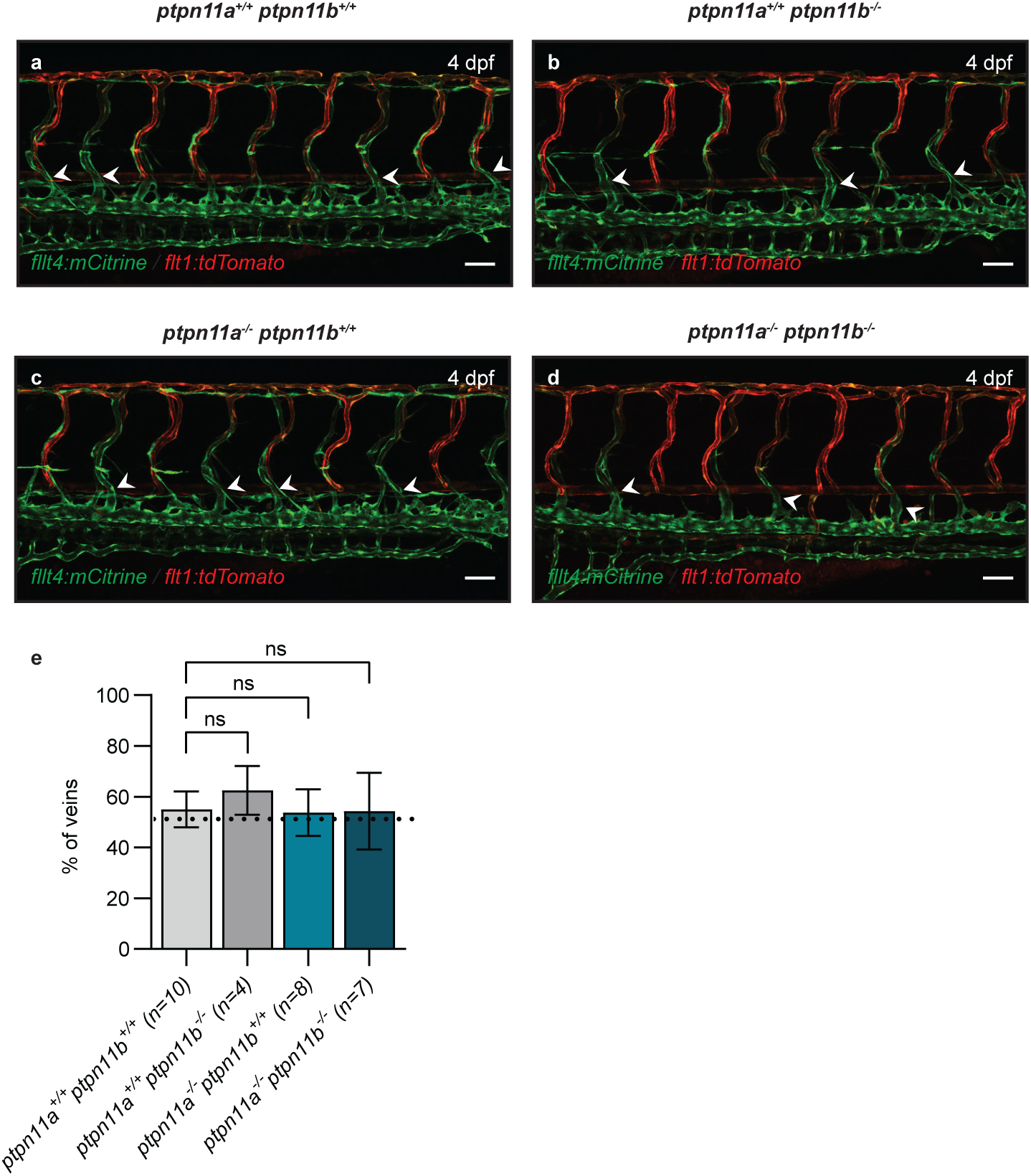
Knockout of Shp2 does not affect establishment of venous ISVs in the trunk. *Ptpn11a^+/−^ptpn11b^−/−^*fish in *Tg(flt4:mCitrine;flt1:tdTomato)* background were incrossed. (**a-d**) Representative confocal projections of the trunk vasculature of single and double homozygous mutant embryos are shown. Scalebar indicates 50 µm. (**e**) Quantification of the percentage of veins in siblings and double homozygous mutants. Statistical analysis was performed using standard t-test. *p < 0.05; **p < 0.01, ***p < 0.001, ****p <0.0001 and ns= not significant.

### *Vegfc* and *Vegfr3* expression is normal prior to secondary sprouting

The VEGFC/VEGFR3 signaling axis is essential for differentiation of endothelial cells to LECs. In the absence of Vegfc or Vegfr3, initiation of lymphangiogenesis is severely impaired (Hogan, Herpers, et al., 2009; Villefranc et al., 2013; Koltowska et al., 2015). SHP2 has a central role in VEGFR signaling. In tissue culture cells, SHP2 has been shown to bind directly to activated VEGFR (Mitola et al., 2006). This results in binding and translocation of the adaptor protein GRB2 to SHP2 and simultaneously, the RAS guanine nucleotide exchange factor, Son of sevenless (SOS), translocates to the cell membrane, resulting in activation of RAS. Hence, loss of Shp2 in zebrafish embryos likely affects Vegfr signaling directly. To assess whether *vegfc* expression is affected by loss of Shp2, we performed comparative *in situ* analysis of *vegfc* at 24, 36 and 48 hpf (**Figure 8, Supplementary Figure 2**). The results show similar expression patterns for siblings and *ptpn11a^−/−^ptpn11b^−/−^*mutant embryos at all stages tested, suggesting that *vegfc* expression was normal (**Figure 8a-c**).

**Figure 8.**
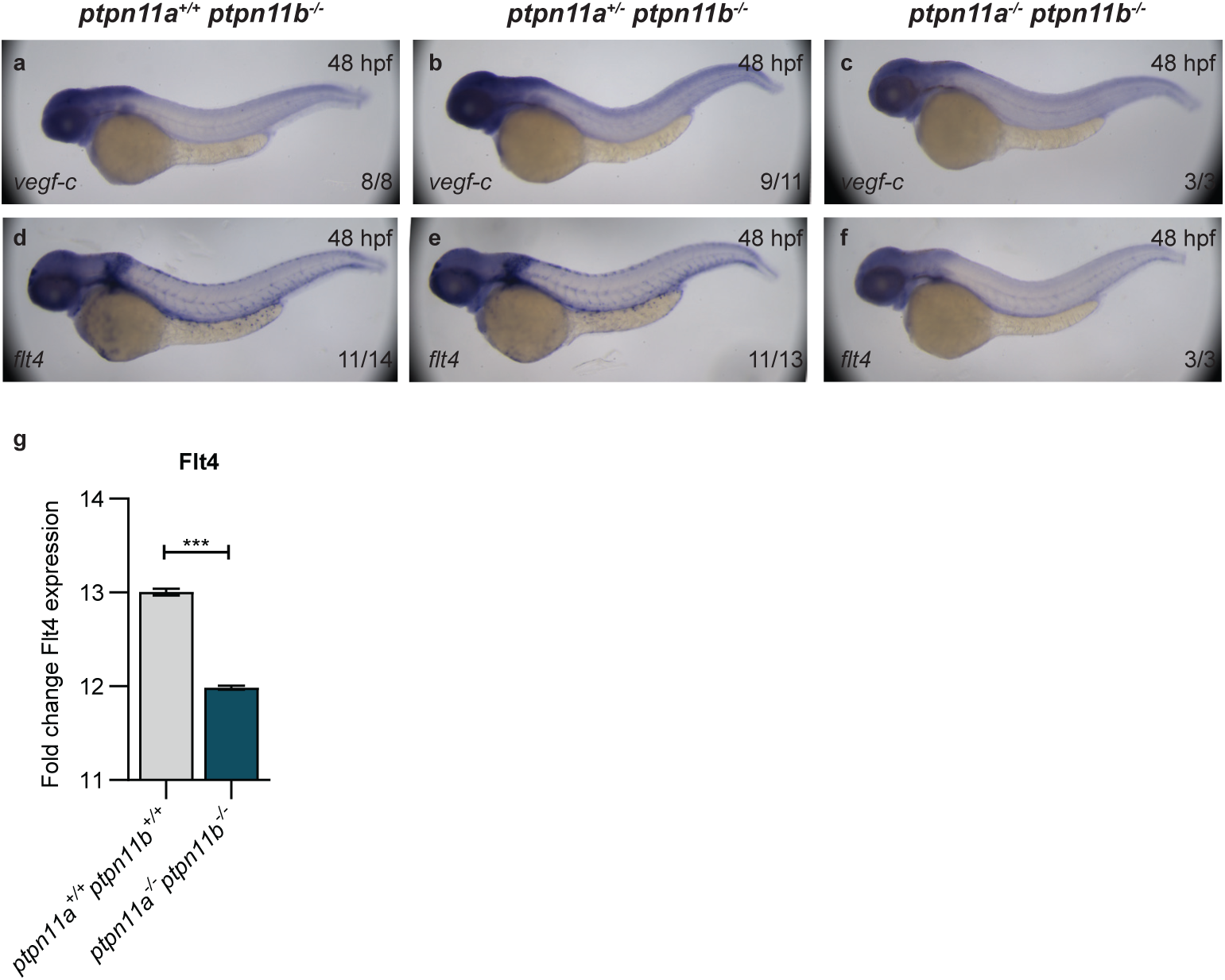
Expression of lymphangiogenic gene *flt4*, but not *vegfc* is downregulated in Shp2 knockout embryos at 48 hpf. *Ptpn11a^+/−^ptpn11b^−/−^*fish were incrossed and embryos were collected at 48 hpf and fixed with paraformaldehyde. *In situ* hybridization was done using (**a-c**) a *vegfc*-specific probe and (**d-f**) a *flt4*-specific probe. In the bottom right corner of each panel the number of embryos showing this pattern of the total number of embryos observed in three independent experiments is depicted. (**g**) Quantitative PCR of *flt4* mRNA expression using cDNA of 5 embryos (5 dpf) per sample. Ct values were normalized using beta-actin and relative expression is shown here (wild-type set to 1.0). Statistical analysis was done using a t-test*, ***p < 0.001*.

Expression of *flt4*, encoding Vegfr3, was similar in all genotypes at 24 and 36 hpf (**Supplementary Figure 2**). However, at 48 hpf when the LEC sprouts are in the process of migrating towards the HM, expression of *flt4* is much less profound in *ptpn11a^−/−^ptpn11b^−/−^*double mutants than in siblings (**Figure 8d-f**). Quantitative PCR indicated a two-fold reduction in *flt4* mRNA expression in *ptpn11a^−/−^ptpn11b^−/−^* embryos compared to wild-type controls (**Figure 8g**), which is consistent with the *in situ* hybridization results. It is noteworthy that activation of VEGFR3 leads to transcriptional activation of *Vegfr3* expression via a positive transcriptional feedback loop in mice (Srinivasan et al., 2014). Our results indicate that expression of *flt4* is significantly reduced in double mutants, which is consistent with Shp2 having a role in downstream Vegfr3 signaling. Moreover, reduced Vegfr3 expression due to impaired Vegfr3 signaling likely amplifies the response to the lack of functional Shp2.

### Phosphatase activity and adaptor function of Shp2 are required for normal lymphangiogenesis

Diverse molecular roles have been identified for the Shp2 protein, including dephosphorylation of downstream signaling molecules as well as serving as an adaptor protein through the N-terminal SH2 domains and the C-terminal tyrosine phosphorylation sites. The catalytic cysteine in the PTP domain (C460 in Shp2) mediates catalytic activity. The absolutely conserved R466 residue is required for catalytic activity and mutation of R466 to methionine renders Shp2 catalytically inactive (Hale & den Hertog, 2018). To address whether lymphangiogenesis depends on the catalytic function of Shp2, we microinjected synthetic mRNA encoding Shp2a variants into one-cell stage embryos obtained from a *ptpn11a^+/−^ptpn11b^−/−^* incross and quantified formation of the TD (**Figure 9a**). The majority of the Shp2a-R466M injected double homozygous *ptpn11a^−/−^ptpn11b^−/−^*embryos lacked the TD and no significant rescue was observed (**Figure 9a**). Morphological analysis indicated that expression of Shp2a-R466M did not rescue the abdominal edema in double homozygous embryos, and the swim bladder remained non-inflated in the microinjected double homozygous embryos (**Supplementary Figure 3**). In *ptpn11a^+/−^ptpn11b^−/−^* and *ptpn11a^+/+^ptpn11b^−/−^*embryos, an increase in the percentage of embryos with a partial or absent TD was observed upon expression of Shp2a-R466M, suggesting a dominant effect of Shp2a-R466M. These results suggest that Shp2 phosphatase activity is required for developmental lymphangiogenesis.

**Figure 9.**
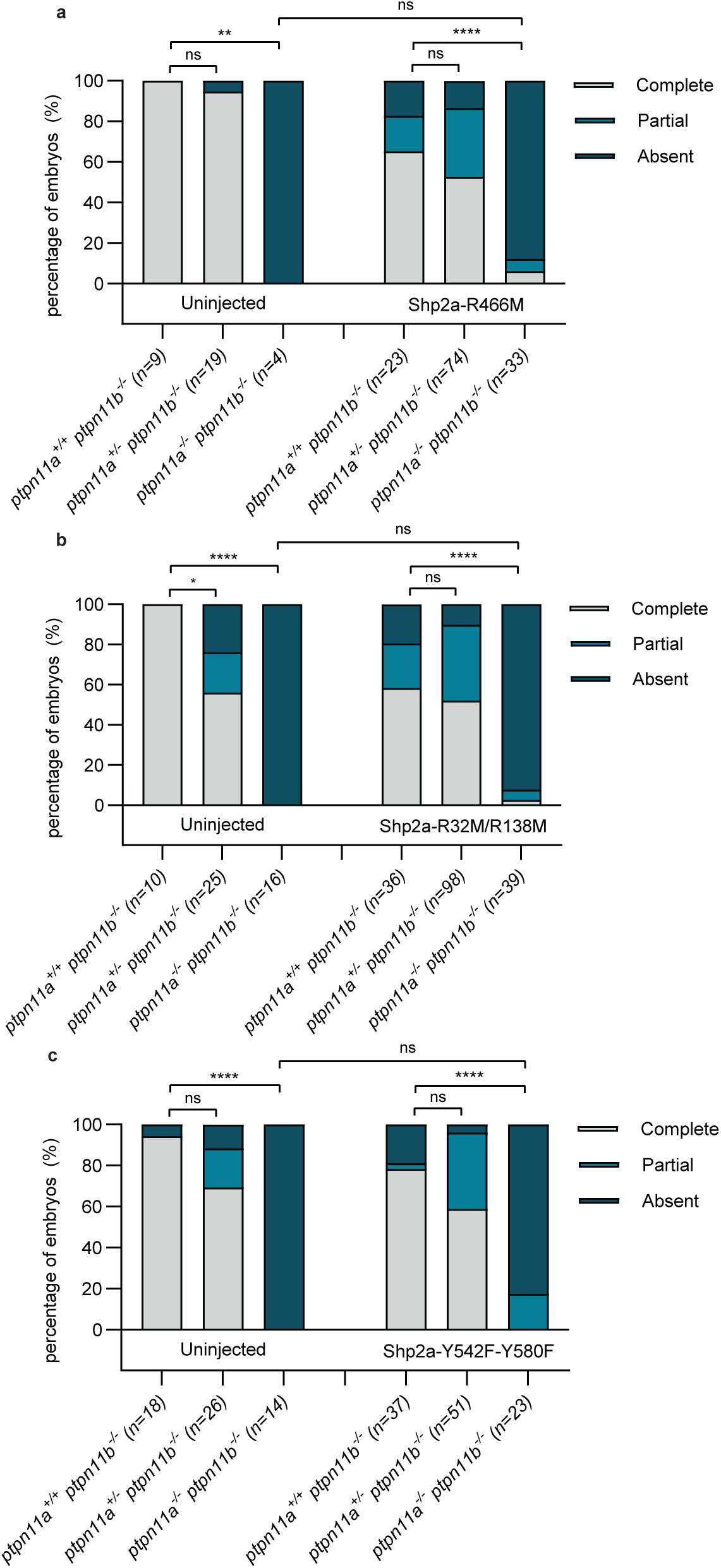
Shp2 catalytic activity, functional SH2 domains and C-terminal Tyr-residues are required for rescue of lymphangiogenesis in Shp2 knockout embryos. *Ptpn11a^+/−^ ptpn11b^−/−^* fish in *Tg(fli1a:eGFP;kdrl:mCherry)* background were incrossed. Embryos were injected at the one-cell stage with synthetic mRNA encoding (**a**) Shp2a-R466M, which lacks catalytic activity, (**b**) Shp2a-R32M/R138M with impaired SH2 domains, or (**c**) Shp2a-Y542F/Y580F, which lacks two phosphorylation sites in the C-terminal tail. At 5 dpf, the presence or (partial) absence of the *fli1a*-positive TD was quantified in uninjected controls and embryos injected with Shp2a mutant by analysis of 10 segments of each embryo. The results of three independent experiments are pooled with the total number of embryos (n) indicated. Statistical analysis was done using the *Mann-Whitney test *p < 0.05; **p < 0.01, ***p < 0.001, ****p <0.0001 and ns= not significant*.

The ‘FLVRES’ motif is conserved in all SH2 domains and encompasses an essential arginyl residue (R32 and R138 in the N-terminal and C-terminal SH2 domain of Shp2, respectively) which is required for binding to pTyr residues in upstream signaling molecules (Eck et al., 1996). Synthetic mRNA encoding Shp2a-R32M/R138M that cannot bind to pTyr-containing proteins through its SH2 domains failed to rescue formation of the TD in *ptpn11a^−/−^ptpn11b^−/−^*embryos (**Figure 9b**). Shp2a-R32M/R138M also did not rescue the morphological defects in *ptpn11a^−/−^ptpn11b^−/−^*embryos, suggesting that functional SH2 domains are required for the function of Shp2 in early lymphangiogenesis (**Supplementary Figure 3**). Expression of Shp2a-R32M/R138M in sibling embryos affected the percentage of embryos with defective lymphangiogenesis (**Figure 9b**).

To establish the role of the Shp2 C-terminal tail for lymphangiogenesis, we microinjected synthetic Shp2a-Y542F/Y580F mRNA into one-cell stage embryos (**Figure 9c**). The characteristic phenotype including severe edema and a non-inflated swim bladder remained evident in the microinjected *ptpn11a^−/−^ptpn11b^−/−^*embryos (**Supplementary Figure 3**). In addition, quantification of the TD revealed no significant difference in these embryos compared to their uninjected siblings. An increase in the percentage of sibling embryos with a partial or absent TD was observed, suggesting that expression of Shp2a-Y542F/Y580F had a dominant effect (**Figure 9c**). These results indicate that the C-terminal tyrosine phosphorylation sites are required for the function of Shp2. Taken together, catalytic activity as well as the adaptor functions of Shp2 are required for the function of Shp2 in development and lymphangiogenesis.

### Patient-associated Shp2 variants rescue to varying extents

Mutations in *PTPN11* are associated with developmental disorders, including Noonan Syndrome (NS) (OMIM 163950) and NS with multiple lentigines (NSML) (OMIM 151100). The mutations causing NS result in enhanced phosphatase activity and enhanced downstream signaling. In contrast, the NSML-associated mutations result in reduced phosphatase activity (Keilhack et al., 2005; Kontaridis et al., 2006; Tartaglia et al., 2006). Yet, NS and NSML have overlapping symptoms. Interestingly, NS is associated with defective lymphangiogenesis (Kleimeier et al., 2022). We wondered whether NS and NSML variants of Shp2 would be able to rescue the defects in lymphangiogenesis in *ptpn11a^−/−^ptpn11b^−/−^* embryos. Synthetic mRNA encoding common variants of NS (D61G and T73I) and of NSML (A462T and T469M) were microinjected at the one-cell stage of embryos obtained from a *ptpn11a^+/−^ptpn11b^−/−^* incross. The morphology was assessed at 5 dpf and formation of the TD was analyzed (**Figure 10**). Interestingly, the NS variants Shp2a-D61G and Shp2a-T73I did not rescue the phenotype observed in *ptpn11a^−/−^ptpn11b^−/−^* embryos to the same extent as wild-type Shp2a (*cf.* **Figures 4 and 10a**). It is noteworthy that Shp2a-T73I rescued the formation of the TD more effectively than Shp2a-D61G, which may reflect differences in catalytic activity or signaling capacity between these two Shp2 variants. Morphologically, Shp2a-T73I rescued double mutant embryos more effectively than Shp2a-D61G as well, in that heart edema and an uninflated swim bladder was observed more frequently in Shp2a-D61G injected double mutant embryos (**Supplementary Figure 4**). The effects of Shp2a-D61G and Shp2a-T73I on *ptpn11a^+/+^ptpn11b^−/−^* and *ptpn11a^+/−^ptpn11b^−/−^* siblings were not significant. Both NSML variants rescued the morphological defects and the lymphangiogenesis defects in *ptpn11a^−/−^ptpn11b^−/−^*embryos to a similar extent as wild-type Shp2a, without significant effects on the siblings (*cf.* **Figure 4, 10b**). Taken together, expression of NS and NSML variants rescued morphological and lymphangiogenesis defects in *ptpn11a^−/−^ptpn11b^−/−^* embryos. Many more NS– and NSML-associated variants of SHP2 have been identified in human patients. Rescue of the lymphatic defects in shp2 double mutants represents a powerful *in vivo* assay for assessing the lymphatic effects of other patient-derived Shp2 variants.

**Figure 10.**
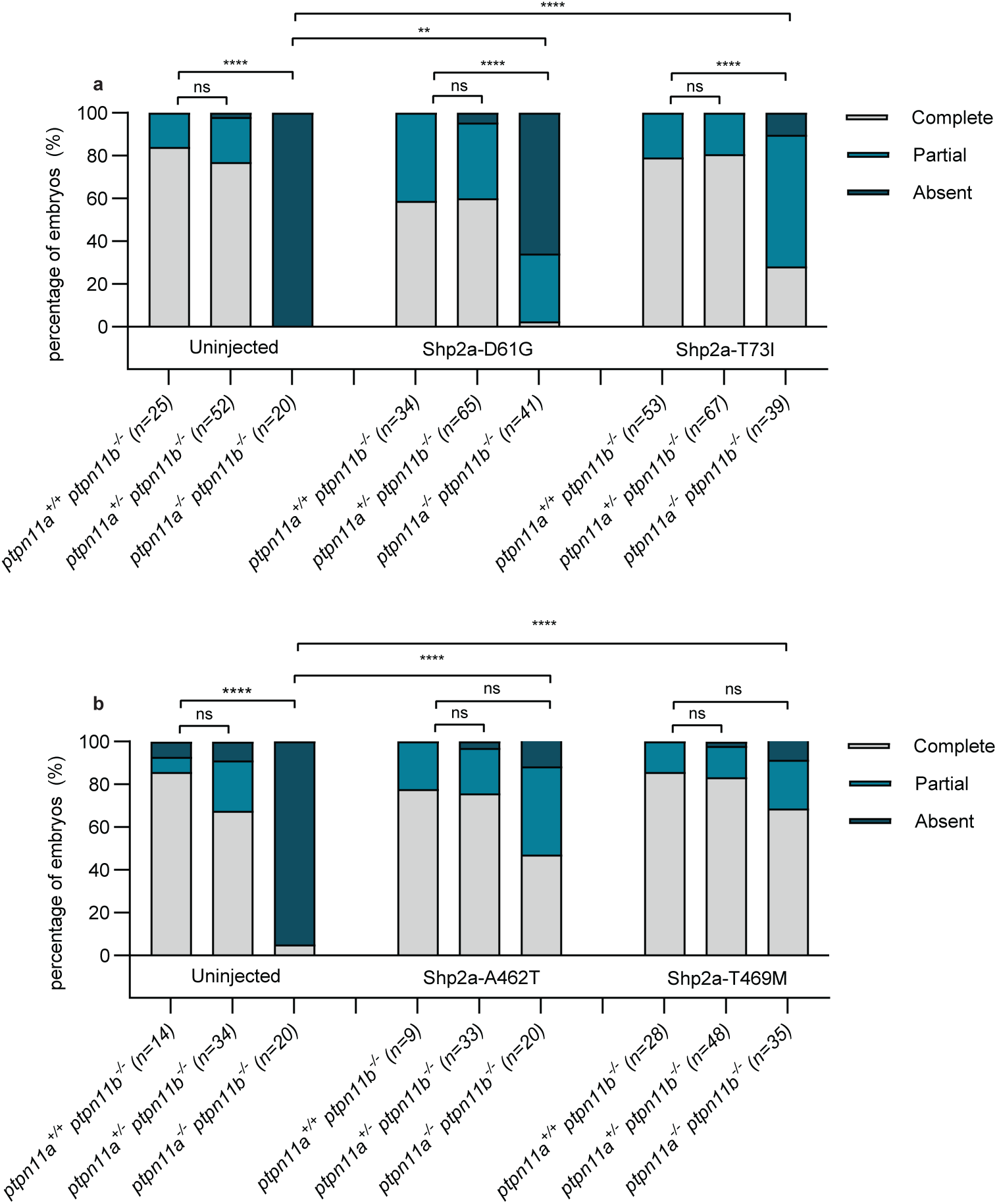
Shp2 NS variants rescue to varying extents and Shp2 NSML variants rescue lymphangiogenesis like wild-type Shp2 in Shp2 knockout embryos. *Ptpn11a^+/−^ptpn11b^−/−^*fish in *Tg(fli1a:eGFP;kdrl:mCherry)* background were incrossed. Embryos were injected at the one-cell stage with synthetic mRNA encoding (**a**) Shp2a NS variants, Shp2a-D61G and Shp2a-T73I, or (**b**) Shp2a NSML variants Shp2a-A462T and Shp2a-T469M. At 5 dpf, the presence or (partial) absence of the *fli1a*-positive TD was quantified in uninjected controls and embryos injected with Shp2a or Shp2b by analysis of 10 segments of each embryo. The results of three independent experiments are pooled with the total number of embryos (n) indicated. Statistical analysis was done using the *Mann-Whitney test *p < 0.05; **p < 0.01, ***p < 0.001, ****p <0.0001 and ns= not significant*.

## Discussion

Lymphangiogenesis is conserved among vertebrates and is driven by the VEGFC/ VEGFR3 pathway (Hogan & Schulte-Merker, 2017). The expansion of available zebrafish models to study this signaling axis has substantially contributed to our knowledge of developmental lymphangiogenesis and exposed key players, including Vegfc, Vegfr3, Ccbe1 and Grb2b (Hogan, Herpers, et al., 2009; Küchler et al., 2006; Mauri et al., 2021; Villefranc et al., 2013). Our research reveals that Shp2, a protein-tyrosine phosphatase with a central role in RAS/MAPK signaling, is required for normal lymphangiogenesis. In embryos that lack functional Shp2, endothelial cell sprouting initiated normally, but cells failed to migrate normally and do not differentiate into PL cells, resulting in a complete lack of PL cells and subsequently of the entire lymphatic vasculature in the trunk of zebrafish embryos.

Facial lymphatics and BLECs were also missing in the absence of Shp2 at 5 dpf. The only exemption was the EC population that gives rise to the “facial collecting lymphatic vessel” (FCLV), which forms the connection between lymphatics and the venous system (Shin et al., 2016). This population is still visible in *vegfc*, *ccbe1* and *flt4* mutants and is therefore considered not to be dependent on Vegfc signaling, but rather on Svep1/Tie1 signaling (Hußmann et al., 2023). Thus, our phenotypic analysis is consistent with the notion that Shp2 acts downstream of Vegfc/ Vegfr3. Interestingly, little or no differences were detected in facial lymphatic structures between wild-type and double mutant at 3 dpf (**Figure 3**), which may reflect the venous origin of the facial lymphatic structures. Whether these facial lymphatic structures subsequently disintegrate for instance by apoptosis in the absence of Shp2 remains to be determined.

Normal formation of lymphatic vessels in the trunk proceeds when endothelial cells of the PCV form protrusions and eventually migrate towards the HM, which is facilitated by local expression and processing of Vegfc (Le Guen et al., 2014; Wang et al., 2020). We confirmed that the gene encoding Vegfc is expressed normally in the double mutant lacking functional Shp2 (**Figure 8**). In addition, time-lapse imaging provided valuable insights into the sprouting behavior of the LEC progenitors in the absence of functional Shp2, suggesting that initial sprouting is unaffected, whereas migration towards the HM is impaired. Normally, the sprouts respond to Vegfc via the VEGFR3 receptor and induce downstream signaling towards the stalk cells to induce proliferation and growth of the sprout (Tammela et al., 2011). This does not occur in embryos lacking functional Shp2, likely because Vegfr3 signaling is impaired due to absence of Shp2.

Venous angiogenesis is not affected by loss of Shp2. There is no difference in the ratio of aISVs:vISVs in double mutants compared to wild-type embryos, suggesting that aISVs are protected from anastomosis with endothelial sprouts by Notch signaling, like in wild-type embryos (Weijts et al., 2018). In double mutant embryos, sprouts developed into vISVs ostensibly without any obstruction. The sprouts protruded apparently normally from the PCV and started to sense the neighborhood with their filopodia (**Supplementary movie 2-3**). Interestingly, all of the sprouts that did not go on to form vISVs in the double mutant embryos stalled and retracted. Subsequently, these cells even went into apoptosis resulting in double mutants without PL cells and subsequently lacking the entire lymphatic vasculature in the trunk.

Defects in lymphangiogenesis are linked, among others, to the VEGFC/VEGFR3 signaling axis, in that knockout of the Vegfc ligand or its receptor, Vegfr3, impairs lymphangiogenesis in all vertebrates. Downstream factors have also been identified to affect lymphangiogenesis, including Grb2b (Mauri et al., 2021). Here, we demonstrate that Shp2 is essential for lymphangiogenesis. SHP2 has a central role in RTK signaling, linking RTKs to GRB2-SOS and further downstream RAS/MAPK signaling (Gale et al., 1993; Rozakis-Adcock et al., 1993). We provide evidence that Vegfr3 signaling is impaired in embryos lacking functional Shp2, in that the positive feedback loop resulting in transcriptional activation of *flt4*, which encodes Vegfr3, is blocked (**Figure 8**). This may actually amplify the effect of loss of Shp2 on Vegfr3 signaling, as reduced levels of Vegfr3 will result in further reduction of signaling that is already reduced due to lack of Shp2.

Interestingly, there are similarities between the previously published *tabula rasa* mutant that lacks functional Grb2b (Mauri et al., 2021) and the Shp2 double knockout. Like the Shp2 double knockout, the Grb2b knockout lacks trunk lymphatic structures and Grb2b has been shown to be indispensable for endothelial cell migration (Mauri et al., 2021). Remarkably, there are also differences in the defects caused by loss of Shp2 and loss of Grb2b. For instance, it is noteworthy that homozygous mutants lacking Grb2b still form PL cells, albeit the number of PL cells is greatly reduced (Mauri et al., 2021), whereas we have not observed any PL cells in the absence of functional Shp2 (**Figure 5**). In addition, there was a significant decrease in the number of vISVs in the *grb2b* mutant compared to wild-type siblings, whereas there was no difference in the Shp2 knockouts. Interestingly, several other zebrafish knockouts have been described with a reduction or complete lack of the lymphatic vasculature, including Svep1 and Trio (Klems et al., 2020; Le Guen et al., 2014). These phenotypes are all accompanied by a decrease in vISVs. The Shp2 double knockout embryos did not show a difference in the percentage of vISVs, underscoring that signaling nuances govern lymphangiogenesis and vISV formation. The Shp2 knockout is the first mutant where LECs are exclusively affected, suggesting that PL cell formation and vISV formation are not strictly coupled processes.

The defects we observed were caused by loss of Shp2, because expression of Shp2 by microinjection of synthetic mRNA at the one-cell stage rescued the defects (**Figure 4**). Multiple signaling functions have been identified in Shp2. The PTP domain exhibits catalytic activity, the N-terminal SH2 domains bind to pTyr-containing proteins and the C-terminal tyrosine phosphorylation sites have been demonstrated to have a role in binding downstream SH2-domain containing proteins. Our results (**Figure 9**) demonstrate that all three signaling functions are required for the function of Shp2 in lymphangiogenesis. Mutation of either function abolished the capacity of Shp2 to rescue the developmental defects in zebrafish Shp2 double mutants. Expression of these three mutants of Shp2 affected TD formation in wild-type and particularly in heterozygous *ptpn11^+/−^ptpn11b^−/−^* embryos, indicating that these Shp2 mutants interfered with endogenous wild-type Shp2 function. Microinjection of NS or NSML variants of Shp2 rescued the developmental defects in Shp2 double mutants to varying extents (**Figure 10**). The two NSML variants (A462T and T469M) rescued developmental defects of Shp2 knockout fish to a similar extent as wild-type Shp2, which is surprising because the NSML variants exhibit very low catalytic activity (Hanna et al., 2006; Jopling et al., 2007; Kontaridis et al., 2006). Catalytically inactive Shp2-R469M did not rescue lymphangiogenesis in Shp2 double knockout embryos (**Figure 9**), suggesting that catalytic activity is required for this function of Shp2. NSML variants of Shp2 have been demonstrated to have some remaining catalytic activity, which may be sufficient for the function of Shp2 (Yu et al., 2013). Alternatively, a phosphatase-independent function of the NSML variants may rescue lymphangiogenesis in Shp2 double knockout embryos. It is noteworthy that neural crest cell survival in response to NSML variants of Shp2 is mediated by a combination of phosphatase-dependent and – independent functions of SHP2 (Stewart et al., 2010). The NS variants rescued lymphangiogenesis in double mutant embryos to varying extents with Shp2-T73I eliciting a rescue that approached the rescue by wild-type Shp2a. Shp2a-D61G, however, rescued poorly. It is noteworthy that Shp2a-D61G affected lymphangiogenesis in siblings of the double mutants as well, which may not be surprising because Shp2a-D61G has a dominant effect over wild-type Shp2. Moreover, defects in the lymphatic vasculature have been observed in human NS patients (Leenders et al., 2025; Pieper et al., 2022). The apparent lack of rescue capacity by the NS variants may be the net result of a partial rescue and simultaneous partial induction of defects in lymphangiogenesis.

Taken together, we have shown that Shp2 is required for the formation of the lymphatic vasculature in zebrafish embryos. Furthermore, we established at which developmental time the Shp2 double mutant phenotype was expressed. The initial sprouts protruded from the PCV but failed to migrate normally to the HM, resulting in a complete lack of PL cells and subsequent lack of lymphatic structures, while vISV formation was unaffected. Our results indicate that Shp2 is dispensable for venous angiogenesis, but is required for lymphangiogenesis, in particular PL cell formation.

## Supporting information

Supplemental Movie 2 part 1

Supplemental Movie 2 part 2

## Acknowledgements

The authors would like to thank the Hubrecht Imaging Center (HIC) for technical assistance and the animal caretakers at the animal facility of the Hubrecht Institute for animal support. The authors are grateful to Bart Weijts (Hubrecht Institute) for invaluable advice. This work was supported by a grant from the DFG (SFB1348 to SSM), an EJPRD grant NSEuroNet (ZonMW 463002003 to JdH) and a KWF Dutch Cancer Society grant (12829 to JdH).

## Notes

### Competing Interest Statement

The authors have declared no competing interest.

