## Supplemental Material for "The protein-tyrosine phosphatase Shp2 is essential for lymphatic endothelial cell differentiation in zebrafish"

### Supplementary Material

**Supplementary movies SM1 and SM2. Aberrant secondary sprouting in double homozygous mutants.** *Ptpn11a*$^{+/-}$*ptpn11b*$^{-/-}$ fish in *Tg(flt4:mCitrine)* background were incrossed and embryos were imaged by time lapse confocal microscopy (1 frame per 5 min). Images were captured using a 20x objective and z-stack step size ranging between 1.5 and 2.0 µm. Cropped confocal projections are shown of a *ptpn11a*$^{+/+}$*ptpn11b*$^{-/-}$ embryo (SM1) and a *ptpn11a*$^{-/-}$*ptpn11b*$^{-/-}$ embryo (SM2 part 1 and part 2). Stills from these movies are shown in Fig. 6.

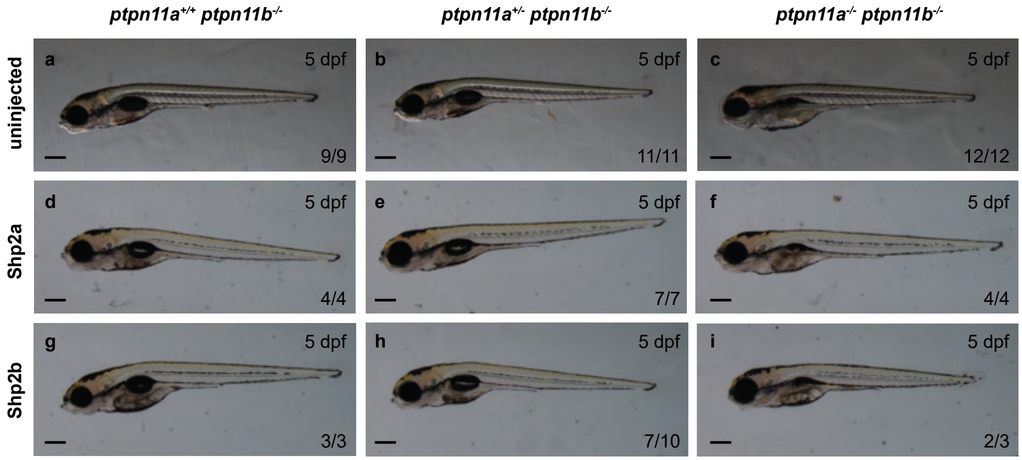


Supplementary Figure 1. Brightfield projections of rescues with Shp2a or Shp2b. *Ptpn11a*$^{+/-}$*ptpn11b*$^{-/-}$ fish in *Tg(fli1a:eGFP;kdrl:mCherry)* background were incrossed. Embryos were injected at the one-cell stage with synthetic mRNA encoding Shp2a or Shp2b. Representative brightfield images of 5 dpf zebrafish embryos in lateral orientation are shown of (a-c) uninjected control, (d-f) Shp2a injected embryos and (g-i) Shp2b injected embryos. Scalebar indicates 300 µm.

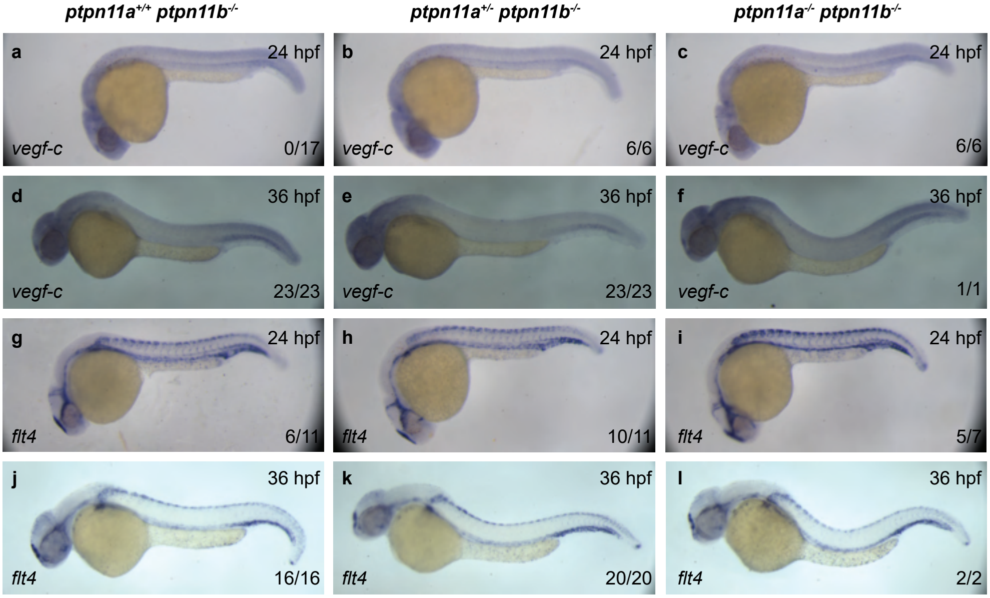


Supplementary Figure 2. Expression of lymphangiogenic genes *vegfc* and *flt4* is not affected in Shp2 knockout embryos at 24 hpf and 36 hpf. *Ptpn11a$^{+/-}$ptpn11b$^{-/-}$* fish were incrossed and embryos were collected at 24 hpf or 36 hpf and fixed with paraformaldehyde. *In situ* hybridization was done using (a-f) a *vegfc*-specific probe and (g-l) a *flt4*-specific probe. In the bottom right corner of each panel the number of embryos showing this pattern of the total number of embryos observed in three independent experiments is depicted.

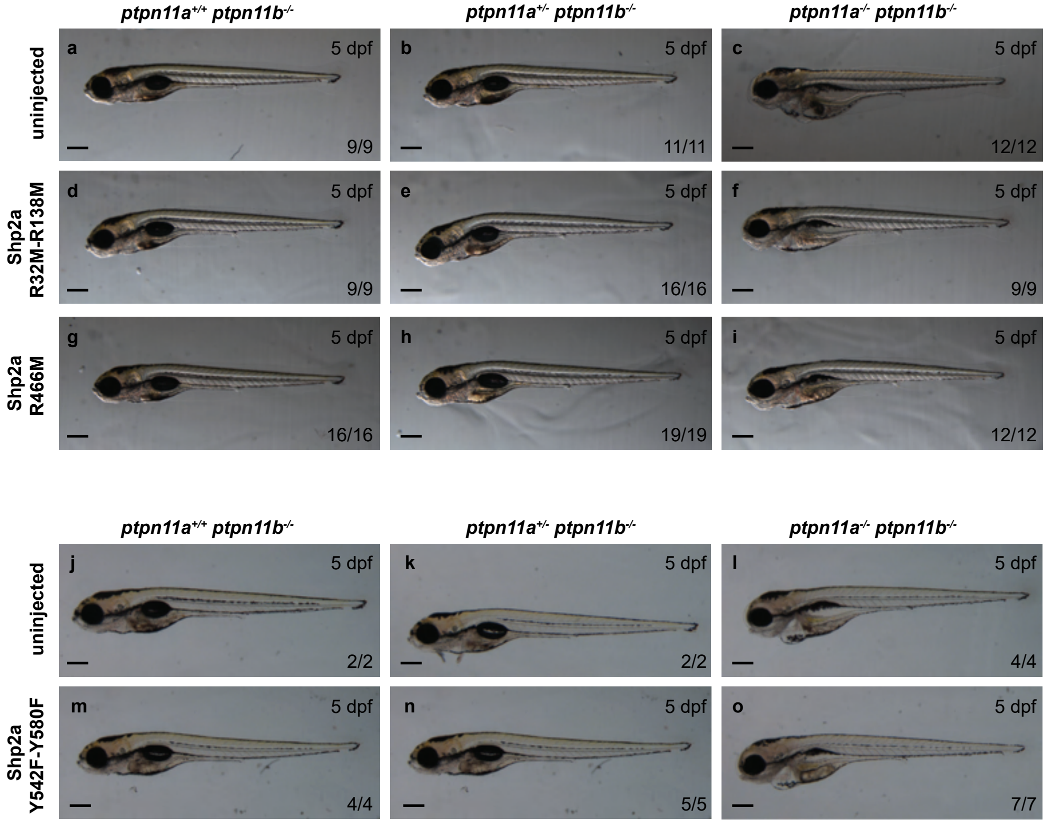


Supplementary Figure 3. Shp2a-R466M, Shp2a-R32M/R138M and Shp2a-Y542F/Y580F did not rescue morphological defects in Shp2 knockout embryos. *$Ptpn11a^{+/-}ptpn11b^{-/-}$* fish in *Tg(fli1a:eGFP;kdrl:mCherry)* background were incrossed. Embryos were not injected (a-c, j-l), or injected at the one-cell stage with synthetic mRNA encoding (d-f) Shp2a-R32M/R138M, (g-i) Shp2a-R466M (j-l), or (m-o) Shp2a-Y542F/Y580F. Representative brightfield images of 5 dpf zebrafish embryos in lateral orientation. Scalebar indicates 300 µm.

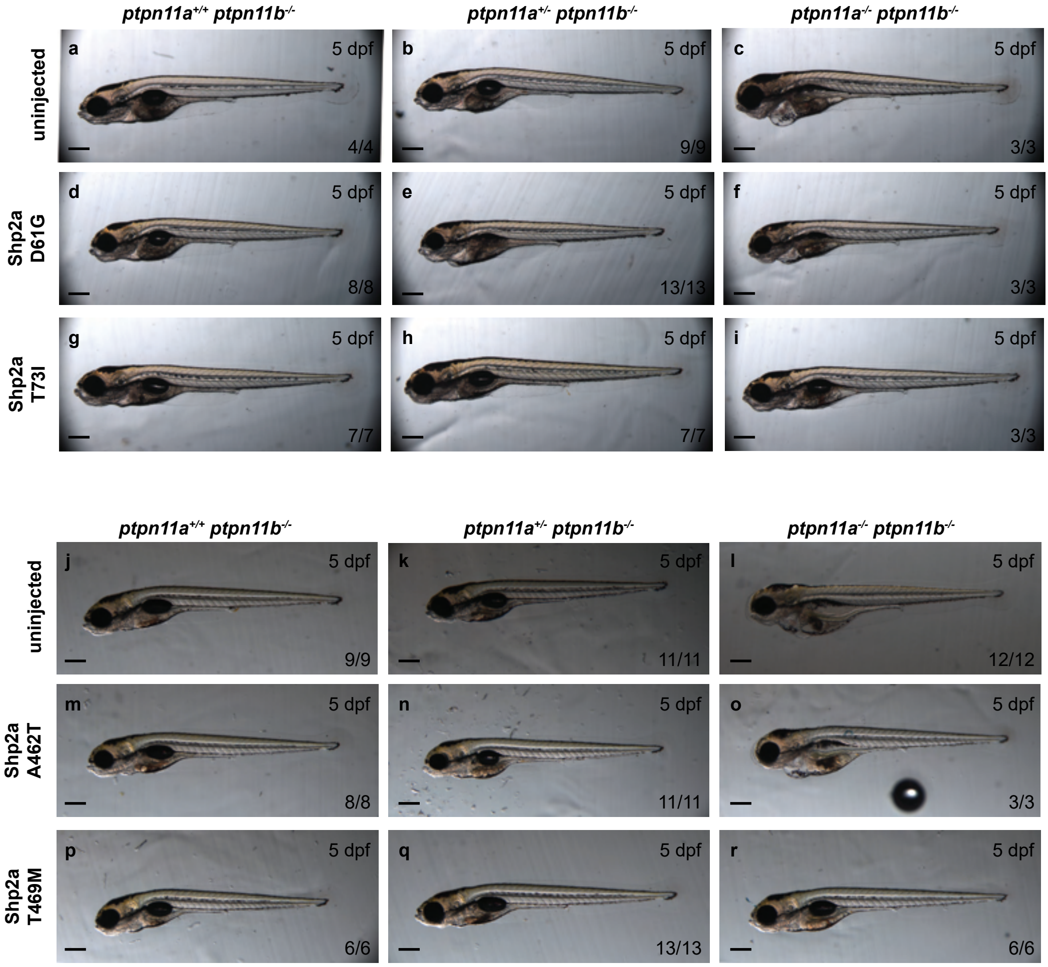


Supplementary Figure 4. NSML variants, but not NS variants rescue morphological defects in Shp2 knockouts. Ptpn11a$^{+/-}$ptpn11b-/- fish in *Tg(fli1a:eGFP;kdrl:mCherry)* background were incrossed. Embryos were not injected (a-c, j-l), or injected at the one-cell stage with synthetic mRNA encoding (d-f) Shp2a-D61G, (g-i) Shp2a-T73I, (m-o) Shp2a-A462T and (p-r) Shp2a-T469M. Representative brightfield images of 5 dpf zebrafish embryos in lateral orientation. Scalebar indicates 300 μm.
